# Cell-autonomous transcriptional mechanism for enhancement of translation capacity in secretory cells

**DOI:** 10.1101/454421

**Authors:** Konstantin Khetchoumian, Aurélio Balsalobre, Alexandre Mayran, Helen Christian, Valérie Chénard, Julie St-Pierre, Jacques Drouin

## Abstract

Translation is a basic cellular process and its capacity is adapted to cell function. In particular, secretory cells achieve high protein synthesis levels without triggering the protein stress response. It is unknown how and when translation capacity is increased during differentiation. Here, we show that the transcription factor Creb3l2 is a scaling factor for translation capacity in secretory cells and that it directly binds ~75% of regulatory and effector genes for translation. In parallel with this cell-autonomous mechanism, implementation of the physiological UPR pathway prevents triggering the protein stress response. The pituitary differentiation factor Tpit activates Creb3l2 expression, the Creb3l2-dependent regulatory network as well as the physiological UPR pathway. Thus, Creb3l2 implements high basal translation levels through direct targeting of translation effector genes acting downstream of signaling pathways that otherwise regulate protein synthesis. Expression of Creb3l2 may be a useful means to enhance production of therapeutic proteins.

---

During embryogenesis, specialized cells acquire different size, shape and organelle content. Some molecular mechanisms at the origin of this morphological diversity were identified in recent years. Indeed, master or scaling transcription factors (TFs), often of the bZIP or bZIP-bHLH type, were associated with biogenesis of endoplasmic reticulum (ER) ^1^, Golgi ^2–4^, mitochondria ^5–7^, lysosomes ^8^ or autophagy ^9,10^ (reviewed in ^11^). In addition, differentiated cells adapt to different physiological, environmental or pathological stresses. The best characterized stress response is the ER stress response, also known as Unfolded Protein Response (UPR). In mammalian cells three ER transmembrane proteins – IRE1, PERK and ATF6 are dealing with ER stress by activating a TF: XBP1s, ATF4 and ATF6N, respectively (reviewed in ^12^). These UPR regulators are the core of a broader family that includes “non-canonical” UPR regulators ^13^, namely the OASIS family of transmembrane bZIP TFs that are ATF6 structural homologues and that have tissue-restricted expression patterns. Their roles remain to be defined in many cases ^14^.

Secretory cells, such as pituitary hormone producing cells, have particularly high protein synthesis requirements in the adult where they function as hormone producing factories. For example, pituitary pro-opiomelamocortin (POMC) secreting cells (corticotropes of the anterior lobe, AL, and melanotropes of the intermediate lobe, IL) increase their hormone production about 100-fold after birth ^15^. The mechanism for this postnatal maturation is unknown. Pituitary hormone production also adapts to physiological or environmental conditions. For example during lactation, prolactin producing cells increase size and expand the Golgi compartment ^16^, while in frogs exposed to dark environment, pituitary melanotropes increase secretory capacity and αMSH production to stimulate skin pigmentation ^17^. The pituitary thus represents an ideal system to study these mechanisms.

Terminal differentiation of POMC-expressing pituitary cells is triggered by Tpit, a Tbox TF only expressed in these cells ^18^. Tpit deficiency results in loss of POMC expression and human *TPIT* mutations cause isolated ACTH deficiency ^19,20^. To identify mechanisms of POMC cell adaptation to the heavy biosynthetic burden happening at the fetal-to-adult transition, we used POMC deficient models to show Tpit-dependent control of translation and secretory capacity through activation of two bZIP TFs, Creb3l2 and XBP1. These TFs exert their cell-autonomous action through direct targeting of genes implicated in translation and ER biogenesis, respectively.

## Results

### Establishment of secretory capacity

As marked upregulation of POMC expression is the hallmark of POMC cell postnatal maturation, we first assessed if this process is dependent on differentiation and/or POMC itself. Inactivation of the *Tpit* gene results in loss of POMC expression in both corticotropes and melanotropes ^20^. In addition, Tpit-deficient pituitaries show a dramatic reduction of intermediate lobe (IL) size (Fig. 1a,b), suggesting there are either fewer cells or decreased cell size. To test the first hypothesis, total IL DNA content was determined. Wild-type (WT) and *Tpit* knock-out (KO) tissues contained the same amount of DNA (Fig. 1c), indicating that cell number is not affected in the absence of Tpit. In contrast, the RNA content of *Tpit* KO IL was reduced 7.9-fold (Fig. 1d). Moreover, IL nuclear staining (Hoechst) showed increased nuclear density in mutant IL (Fig. 1a,b, insets), suggesting that Tpit-deficient cells are smaller. FACS analysis confirmed this, and also revealed reduced organelle content (granularity) (Fig. 1e,f). The reduction of *Tpit* KO melanotrope cell volume was found to be 7-fold compared to WT (Fig. 1g), while cell granularity was decreased 3-fold (Fig. 1h). Thus, postnatal maturation of secretory organelle content appears to be Tpit-dependent.

**Figure 1.**
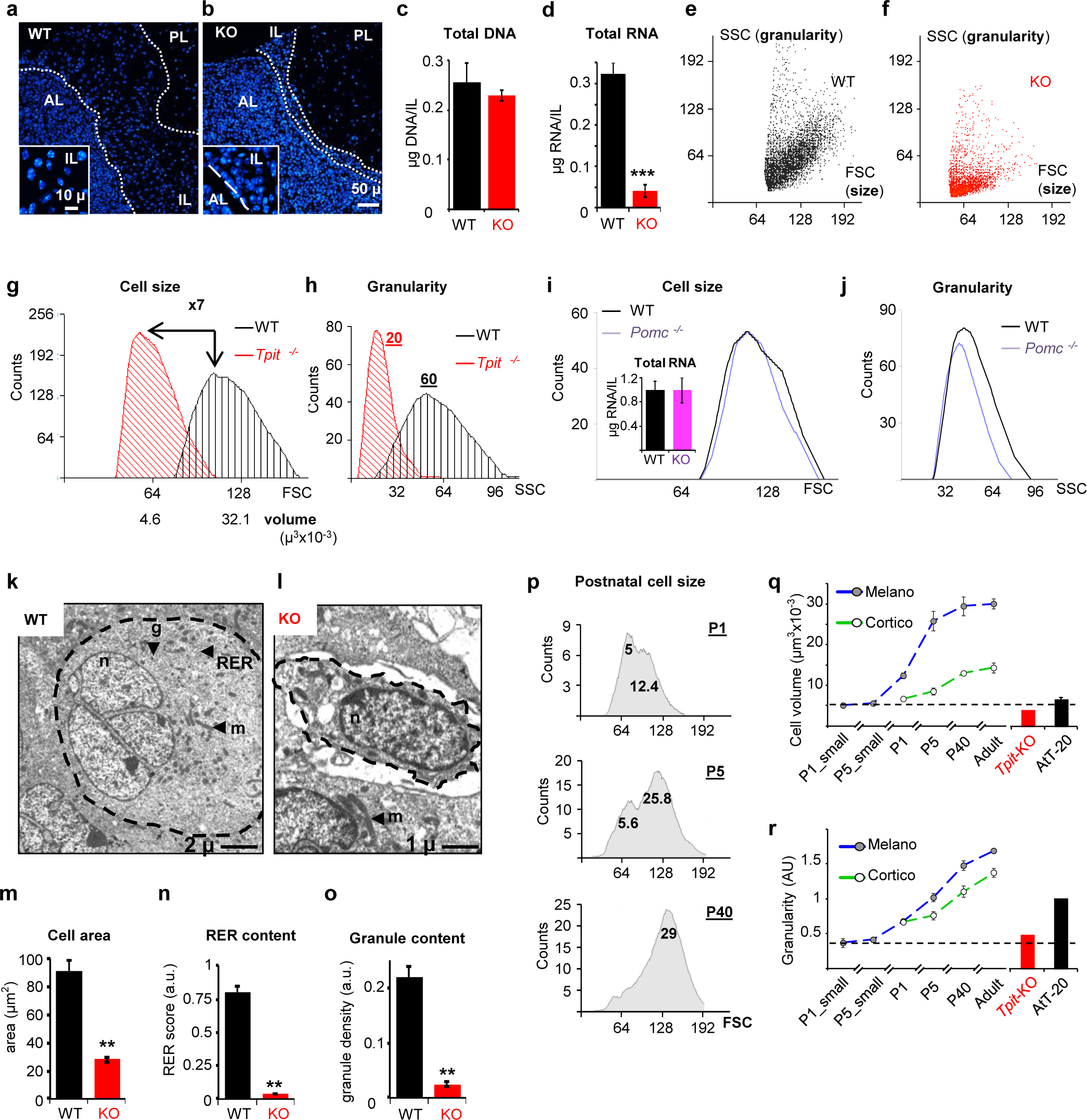
Tpit is required for postnatal maturation of pituitary POMC cells. (**a-o**) Reduced cell size and organelle content in Tpit-deficient pituitaries. (**a,b**) Nuclear staining (Hoechst) of pituitary sections from adult WT (**a**) and KO (*Tpit*−/−, **b**) mice. Demarcations between pituitary lobes (anterior: AL, intermediate: IL, posterior: PL) are indicated by white dashed lines. Higher magnification insets show increased nuclear density in mutant IL. (**c**) Quantitation of IL total genomic DNA contents indicates that cell number is not affected in *Tpit* KO. (**d**) Total IL RNA content is reduced in *Tpit* KO tissues. (**e-h**) Flow cytometry analysis of WT (black) and *Tpit-KO* (red) IL cells showing forward (FSC) *vs* side (SSC) scatter analyses that reflect cell size and granularity (organelle content), respectively. Representative FACS profiles are shown for WT (**e**) and KO (**f**) IL. Graphs show distribution of cell size (**g**) and granularity (**h**) for each genotype. (**i,j**) RNA content (histogram) and FACS analyses of *Pomc*-deficient IL cells (purple) indicate normal RNA content, size (**i**) and slightly reduced granularity (**j**). (**k-o**) Ultrastructure analyses confirm cell size and organelle content defects in *Tpit*-deficient IL. (**k,l**) Electron micrographs of sections from WT (**k**) or KO (**l**) adult mouse pituitaries. WT cells (**k**) are rounded (dashed line), contain dense-cored secretory granules (g), mitochondria (m) and some rough endoplasmic reticulum (RER). Mutant cells (**l**) are smaller, with little cytoplasm and organelles (mitochondria dominate) and have a stellate appearance (not rounded). (**m-o**) Quantitation of cell area (**m**), RER content (**n**) and granule density (**o**) in WT or KO IL cells. Data are presented as means (n=4 mice) ±SEM. Scale bar in **k**=2 µm, in **l**=1 µm. n=nucleus. (**p-r**) Pituitary POMC cells develop into secretory factories postnatally. (**p**) Time-course of melanotrope cell size changes after birth. FACS profiles showing FSC (forward scatter as measure of size) distribution of GFP-positive cells of intermediate lobe (IL) melanotropes from *POMC-EGFP* transgenic mice. Numbers under the peaks indicate calculated cell volumes (µm^3^×10^−3^). (**q,r**) Summary of size (**q**) and granularity/organelle content (**r**) changes in postnatal IL melanotropes (filled circles) and AL corticotropes (empty circles). Inferred progression of cell size (FSC) and granularity (derived from side scatter, SSC) in melanotropes (blue) and corticotropes (green) between days P1 and P90 (adult) are shown by dashed lines. Size and granularity of *Tpit-KO* cells remain at the P1 stage (red bars) and AtT-20 cells appear as weakly secreting (immature) corticotropes (black bars).

The decrease of cell size and granularity may be secondary to the absence of POMC mRNA in *Tpit* KO cells since this mRNA constitutes their major translation burden. We used *Pomc* KO IL cells to assess this possibility. Strikingly, IL RNA content, cell size and organelle contents were not affected by the absence of POMC mRNA (Fig. 1i,j). In order to directly ascertain the putative loss of organelles in Tpit-deficient cells, we performed electron microscopy. Whereas WT melanotropes (Fig. 1k) are rounded, contain dense secretory granules, mitochondria and rough endoplasmic reticulum (RER), KO cells (Fig. 1l) appeared to be smaller, with little cytoplasm or organelles. Quantitation of these features revealed reduced cell area, RER and granule content (Fig. 1m-o) in KO IL cells. In summary, postnatal maturation of pituitary POMC cells is part of the Tpit-dependent differentiation program and is not secondary to the translational burden of the POMC mRNA.

In addition to the 100-fold increase of POMC mRNA levels in adults^15^, examination of POMC cells suggested that they increase in volume during postnatal development. We took advantage of *POMC-EGFP* reporter mice^15^ to analyze by FACS the time course of this increase. Both melanotropes and corticotropes increase in size between postnatal days P1 and P40, with greater amplitude in melanotropes (Fig. 1p, q). In addition, an increase of cell granularity was observed (Fig. 1r), suggesting an expansion of organelle content. In summary, maturation of POMC cell secretory capacity is implemented during the postnatal period and it is triggered by Tpit.

### Creb3l2, a Tpit-dependent regulator

To gain insights into the molecular mechanisms of Tpit-dependent POMC cell maturation, we compared gene expression profiles of WT and *Tpit* KO IL that contain mostly melanotropes ^21^. Comparison of WT *versus* KO gene expression profiles (Supplementary Fig. S1a-c) revealed 2697 differentially-expressed transcripts using a *P*-value cut-off *P*<0.001 (Fig. 2a and Supplementary Table S1). Gene Ontology (GO) analysis revealed that the most significantly enriched biological processes (BP) associated with 1578 transcripts downregulated in Tpit-deficient IL are intracellular protein transport, secretory pathway and translation (Fig. 2b). In particular and as validated by RT-qPCR, major regulators of the UPR pathway – XBP1, ATF4 and ATF6, as well as numerous downstream UPR pathway, vesicle-mediated transport and translation control genes are Tpit-dependent (Fig. 2c,d). These genes were not downregulated in *Pomc*-KO ILs (Fig. 2d). Hence, expression of Tpit, but not POMC, correlates positively with that of secretory pathway genes. Interestingly, UPR genes associated with translational repression (eIF2α kinase *Perk/Eif2ak3*), apoptosis (*Chop/Ddit3*) or ER associated protein degradation (ERAD) are not affected by loss of Tpit (Supplementary Fig. S1d). In addition, expression of *Ppp1r15a*/*Gadd34/Myd116* (phosphatase that reverses inhibitory eIF2α phosphorylation) and *Naip5*-*Naip6* (anti-apoptotic genes) is decreased in *Tpit KO* IL. Therefore, Tpit action correlates with activation of some branches of the UPR pathways but not with those involved in translational attenuation, disposal of misfolded proteins and programmed cell death that are part of the classical XBP1-dependent UPR stress response. These differences distinguish the physiological UPR from the ER stress-related UPR response.

**Figure 2.**
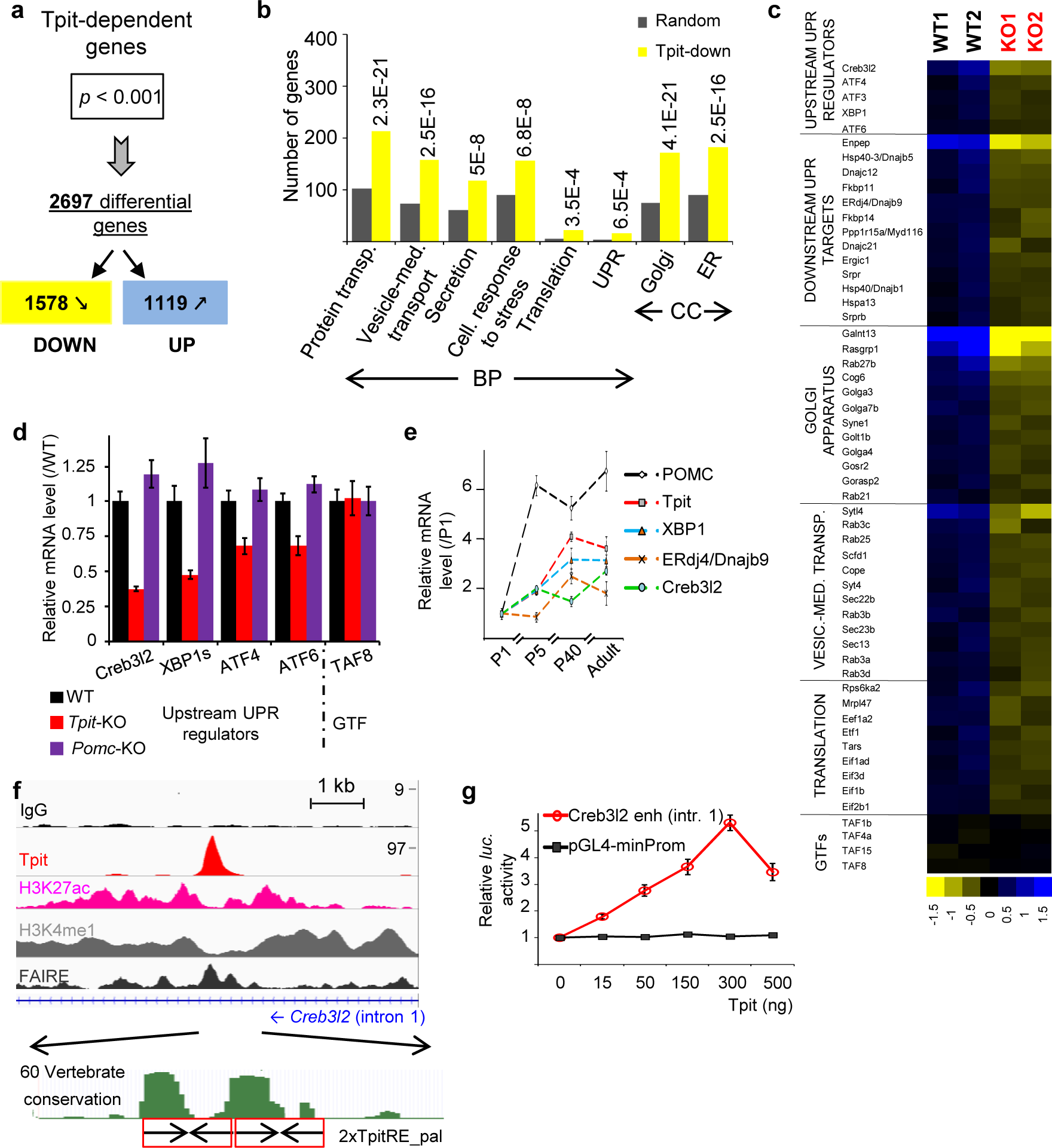
Tpit controls expression of translation and secretory pathway genes. (**a-e**) Tpit transcriptional signature in the pituitary. (**a**) Tpit-dependent genes identified by comparison of WT and KO IL transcriptomes. (**b**) Gene ontology (GO) distribution of transcripts downregulated in *Tpit*-deficient IL showing enrichment of protein transport, secretory pathway and translation genes. BP, biological process. CC, cellular component. Bars represent the number of genes downregulated in *Tpit*-deficient ILs (yellow) and random occurrence of genes in each category (grey). *P*-values relative to random occurrence are shown above the bars. (**c**) Heat map representation (log_2_ of changes relative to median) of expression for a subset of genes in two WT and two KO IL samples illustrating reduced level of UPR/secretory pathway/translation genes in *Tpit*-deficient tissues. Each column represents an IL sample, and each row represents a gene. Blue and yellow represent up- and down-regulation, respectively. Expression levels of general transcription factors (GTFs) are not affected by *Tpit* KO. (**d**) Real-time quantitative PCR (RT-qPCR) validation of expression changes for a subset of UPR pathway genes observed in *Tpit*-deficient tissues (red) and in *Pomc*-deficient ILs (purple). For each gene, relative mRNA levels were normalized to TBP mRNA and represented as fraction of WT expression levels (black, adjusted to 1). Data represent averages (±SEM) of 3-6 mice per genotype. (**e**) Postnatal expression of *Tpit, Pomc, Creb3l2* and UPR pathway genes in melanotropes. For each gene, relative IL mRNA levels measured by RT-qPCR were normalized to those of TBP and represented as fraction of P1 expression level (set to 1). Data represent averages (±SEM) of 5-8 animals per group. (**f,g**) Creb3l2 is a major Tpit target. (**f**) *Top*: ChIPseq profiles in AtT-20 cells revealing direct recruitment of Tpit to a *Creb3l2* intron 1 sequence exhibiting active enhancer marks, namely bimodal H3K4me1, H3K27ac and FAIRE peaks. *Bottom*: two conserved Tpit-RE palindromic sequences at the Tpit recruitment region of *Creb3l2* intron 1. (**g**) Dose-dependent activation of a *Luciferase* reporter containing the intron1 enhancer (986 bp) of the *Creb3l2* gene by Tpit assessed by transfection into αT3 pituitary cells. The control plasmid (pGL4) contains the minimal -34 bp *Pomc* promoter only.

The non-canonical bZIP TF Creb3l2 is the most downregulated gene of this subfamily in *Tpit* KO (Fig. 2c,d and Supplementary Fig. S1e) and it is noteworthy that the pituitary is a major tissue of Creb3l2 expression (Supplementary Fig. S1f). Expression of both *Creb3l2* and *XBP1*, the classical UPR responsive TF, follow the Tpit/POMC postnatal expression profile (Fig. 2e), in agreement with the hypothesis that they mediate Tpit actions on postnatal maturation of POMC cells. In order to correlate Tpit-dependence with Tpit action, we queried the Tpit ChIPseq data ^22^ for direct binding to putative *Creb3l2* regulatory sequences. Strong Tpit binding (8^th^ highest peak out of 17190) is present in the first intron of the *Creb3l2* gene (Fig. 2f). This site contains two conserved juxtaposed TpitRE palindromic sequences ^23^ and exhibits the chromatin signature (H3K4me1, H3K27ac and FAIREseq peak) of active enhancer sequences. When directly tested in a luciferase reporter assay, this sequence behaved as a *bona fide* Tpit-responsive enhancer (Fig. 2g). Altogether, these results identify *Creb3l2* as a major Tpit target gene.

### Translation and secretory capacity

In order to evaluate *in vivo* the putative roles of Creb3l2 and XBP1, we performed loss- and gain- of function (LOF, GOF) experiments. Because Creb3l2 or XBP1 germline inactivation lead to perinatal or embryonic lethality ^24,25^, we used a pituitary POMC cell specific dominant-negative (DN) approach. Replacement of the basic region of bZIP proteins by an acidic (A) sequence (AZIP) was shown to be a very efficient and specific way to inactivate endogenous bZIP factors^26^ (Fig. 3a). We generated ACreb3l2 and AXBP1 constructs and validated their efficiency and specificity using a 3xCreb3l2-RE-luc reporter in both activated and basal conditions (Fig. 3b and Supplementary Fig. S2a, b). To inhibit Creb3l2 or XBP1 in pituitary POMC cells, we expressed ACreb3l2 or AXBP1 in transgenic mice using the POMC promoter, previously shown to be specific and penetrant in melanotrope cells ^15^. For each transgene, we selected two stable mouse lines with an AZIP versus endogenous bZIP mRNA ratios >5, ensuring strong inhibition of the endogenous TF (Fig. 3b and Supplementary Fig. S2c, d). Both transgenic lines behaved similarly and results are shown for one line. Total RNA content of transgenic ILs was reduced with synergistic effects in double ACreb3l2/AXBP1 transgenic mice (Fig. 3c). Also, ACreb3l2 and AXBP1 increased IL nuclear density, reflecting a reduction in melanotrope cell size (Fig. 3d, e).

**Figure 3.**
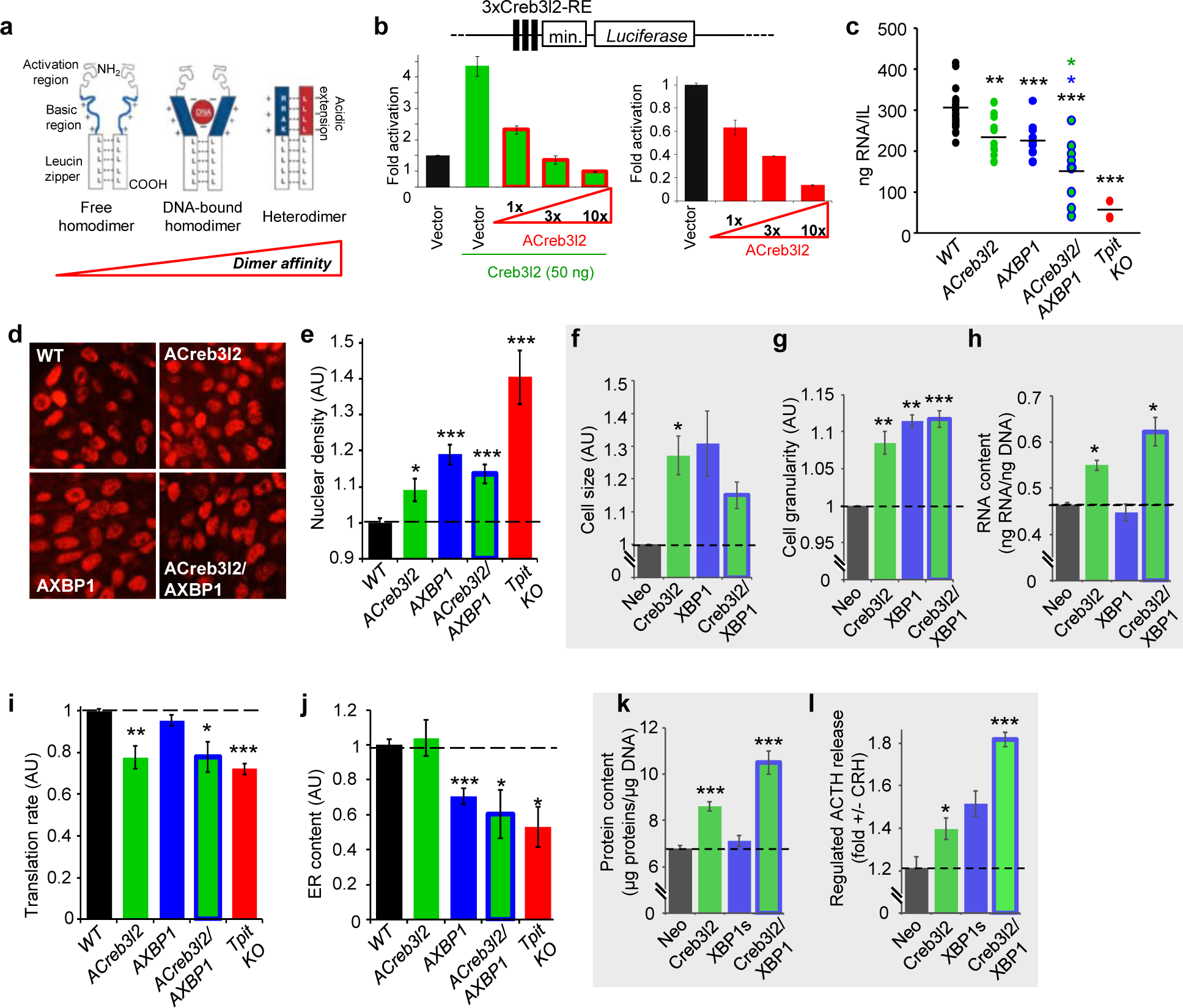
Creb3l2 and XBP1 regulate different aspects of secretory capacity. (**a**) Schematic view of dominant negative inhibition of bZIP TFs by AZIP proteins (adapted from ^26^). *Left*: free bZIP/bZIP homodimer with unstructured basic region (blue). *Middle*: DNA bound bZIP/bZIP homodimer with α-helical basic region. *Right*: bZIP/AZIP heterodimer where the basic region and the designed acidic amphipathic extension (red) interact as α-helices to extend the coiled-coil domain and prevent DNA binding. (**b**) Efficiency of overexpressed (*left*) or endogenous (*right*) Creb3l2 inhibition by ACreb3l2. The Creb3l2-RE *Luciferase* reporter (3xCreb3l2-RE-*luc*) was used to assess Creb3l2 activity and ACreb3l2 antagonism upon transfection into INS-1 cells. (**c**) Total IL RNA contents measured in WT, *ACreb3l2*, *AXBP1*, *ACreb3l2/AXBP1* transgenic and *Tpit-KO* mice. (**d,e**) Melanotrope nuclear density (Tpit nuclear staining: red) on histological sections of ILs from WT, *ACreb3l2*, *AXBP1* and *ACreb3l2/AXBP1* transgenic mice (**d**) and their quantifications in arbitrary units (AU) relative to WT melanotrope density adjusted to 1 (**e**). (**f-h**) Creb3l2 and XBP1 gain of function in stably infected AtT-20 cells. FACS analyses showing cell size (**f**) and granularity (**g**) index for AtT-20 cells expressing Creb3l2, XBP1 or both compared to control neomycin-resistant (Neo) AtT-20 cells. Data (±SEM, n=3) are presented relative to Neo cells set to 1. (**h**) Total RNA content of control AtT-20 cells (Neo) and AtT-20 cells expressing Creb3l2, XBP1 or both (±SEM, n=3). (**i-l**) Relative contributions of Creb3l2 and XBP1 to secretory capacity. (**i**) SUnSET (puromycin incorporation) measurements of protein synthesis in IL cells of control (WT), *ACreb3l2*, *AXBP1*, *ACreb3l2/AXBP1* transgenic and *Tpit-KO* mice. Data in AU are presented as fractions of WT IL translation rates set to 1. (**j**) Melanotrope cell ER contents (ERtracker) measured in WT, *ACreb3l2*, *AXBP1*, *ACreb3l2/AXBP1* transgenic and *Tpit-KO* mice. Data are presented as fraction of WT IL ER contents set to 1. (**k**) Total protein content of control AtT-20 cells (Neo) and AtT-20 cells expressing Creb3l2, XBP1 or both. (**l**) CRH-induced ACTH release from the indicated pools of AtT-20 cells.

To complement these *in vivo* LOF models, we assessed the consequences of Creb3l2 or/and XBP1 overexpression (GOF) on size and secretory capacity of POMC cells by generating stable populations of AtT-20 cells that overexpress active forms of Creb3l2 (cleaved Creb3l2) and/or XBP1 (spliced XBP1). Expression of Creb3l2, XBP1 or both led to a similar increase of AtT-20 cell size and granularity (Fig. 3f, g). In contrast, Creb3l2, but not XBP1, overexpression resulted in higher RNA contents, and this effect is even greater in Creb3l2+XBP1 overexpressing cells (Fig. 3h). Hence, Creb3l2 and XBP1 are controlling overlapping (cell size and organelles) and unique (RNA content) aspects of secretory capacity.

As the Tpit-dependent transcriptome (Fig. 2b, c) identified translation and ER biogenesis, we assessed these functions in the *in vivo* LOF and AtT-20 GOF models. Strikingly, Creb3l2, but not XBP1, inhibition led to a specific reduction of translation rate (Fig. 3I), as assessed by the puromycin incorporation assay ^27^. This effect fully mimics the loss of Tpit. In contrast, ER content is dependent on XBP1, but not Creb3l2 (Fig. 3j), and XBP1 inhibition also mimics the *Tpit* KO phenotype. Altogether, these results show that Creb3l2 and XBP1 have different activities, namely regulation of translation and ER biogenesis, respectively. Consistent with the role of Creb3l2 in translation, Creb3l2, but not XBP1 overexpressing cells have higher total protein content (Fig. 3k). An even greater increase was observed in Creb3l2+XBP1 overexpressing cells.

We also assessed the impact of Creb3l2 and XBP1 on the regulated secretory pathway ^28^ by measuring ACTH release in response to corticotropin-releasing hormone (CRH). Both, Creb3l2 and XBP1 overexpression improved CRH response, with a synergistic effect in Creb3l2+XBP1 cells (Fig. 3l). Collectively, these results suggest that Creb3l2 and XBP1 are jointly regulating the development of secretory function.

The Akt/ mTOR pathway regulates translation in response to growth factors; in order to assess whether this pathway is involved in the action of Creb3l2, we measured by Western blot the levels of phospho-mTOR and mTOR. These are largely unaltered by *Tpit* KO, the *ACreb* or *AXBP1* transgenes (Supplementary Fig. S2e) suggesting that Creb3l2 may act downstream of this pathway.

### Creb3l2 targets the promoters of translation genes

In order to define the mechanism by which Creb3l2 may enhance translation capacity and to document XBP1 action on the physiological UPR pathway, we performed ChIPseq for both factors in AtT-20 cells. We observed 6484 Creb3l2 recruitment peaks, while XBP1 ChIPseq produced fewer peaks (258) similar to observations made in other cells ^29,30^. Contrary to many TFs such as Tpit that are mostly recruited at enhancers, 41% of Creb3l2 and XBP1 peaks are found at promoters (Supplementary Fig. S3a-c). While XBP1 targets genes involved in ER biogenesis (Supplementary Fig. S3d) in agreement with previous reports ^29^, Creb3l2 peaks are enriched at regulatory sequences of genes controlling translation (Fig. 4a-e and Supplementary Fig. S3e-h). The bulk of Creb3l2 and XBP1 peaks are associated with active chromatin marks, namely DNA accessibility revealed by FAIRE (“formaldehyde-assisted isolation of regulatory elements”) and methylated H3K4me (Supplementary Fig. S3i, j).

**Figure 4.**
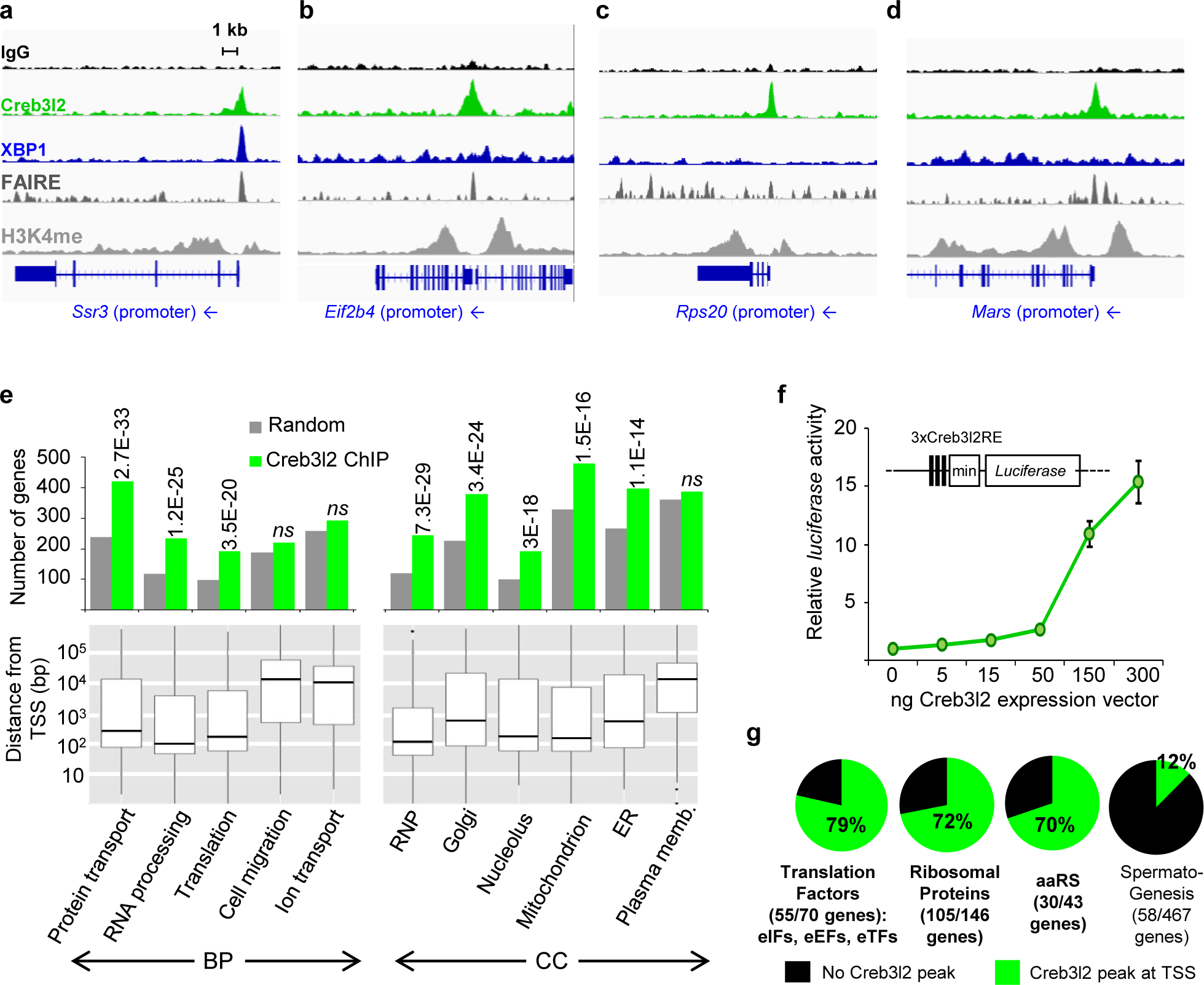
Creb3l2 targets the promoters of translation genes. (**a-d**) ChIPseq profiles for Creb3l2 and XBP1 at regulatory sequences of ER (**a**) and translation (**b-d**) genes. ChIPseq patterns are shown for control IgG, Creb3l2, XBP1, H3K4me1 and FAIRE-seq. (**e**) GO terms of genes associated with TSS proximal (≤1 kb) Creb3l2 peaks. Peaks were assigned to the closest gene with the AnnotatePeaks Homer command. Bars represent the number of genes in each category associated with Creb3l2 peaks (green) or random occurrence of genes in each category (grey). *P*-values relative to random occurrence are shown above the bars. *ns*, not significant. The bottom panels provide box-plot representation of the distance to TSS for Creb3l2 peaks of each GO category revealing promoter-proximal associations for groups with significant associations. (**f**) Dose-dependent activation by Creb3l2 of luciferase reporter containing three copies of Creb3l2 binding sites (3xCreb3l2-RE) upon transfection into pituitary GH3 cells. (**g**) Creb3l2 binds 70-80% of the promoters of translation genes. Pie charts showing proportion of genes (absolute gene numbers indicated between parentheses and target genes listed in Supplementary Table S2) bound by Creb3l2 (green) in each category related to translation, namely: translation factors, ribosomal proteins and aminoacyl tRNA synthetases. The spermatogenesis group serves as control. Gene lists for each category are from MGI (http://www.informatics.jax.org/mgihome/GO/project.shtml).

GO analysis of genes associated with Creb3l2 peaks confirmed enrichment of categories related to translation, protein transport, and RNA processing (Fig. 4e), while XBP1-associated genes correspond to ER stress/UPR (Supplementary Fig. S3d). These associations mostly occur at promoters/proximal regions of target genes (Fig. 4e and Supplementary Fig. S3d, bottom panels). Collectively, these data show targeting of two subsets of genes involved in secretory function, namely genes involved in translation and protein transport (controlled by Creb3l2) and ER biogenesis (controlled by XBP1).

Analysis of the ChIPseq data for *de novo* motifs revealed conserved sequence motifs for Creb3l2 and XBP1 (Supplementary Fig. S4a, b) that are consistent with prior work ^22,23,29,31^. Creb3l2 and XBP1 peaks were associated with bZIP motifs containing the core ACGT sequence. A reporter containing three copies of the Creb3l2-RE upstream of a minimal promoter as well as a natural promoter (*Ssr3*) and enhancer (*Kcnma1*) were activated dose-dependently by both Creb3l2 and XBP1 (Fig. 4f and Supplementary S4c, d). Thus, Creb3l2 and/or XBP1 directly activate genes involved in translation and secretory system development.

In order to better assess the extent of Creb3l2 targeting of genes involved in translation, we computed the number of genes encoding translation initiation (eIF), elongation (eEF) and termination (eTF) factors as well as ribosomal protein coding genes and those for aminoacyl tRNA synthethases that are directly targeted by Creb3l2 binding (within 100 bp of their TSS) based on the ChIPseq data. This showed direct targeting of 70-80 % of these genes depending of the sub-categories (Fig. 4g, Supplementary Table S2). Collectively, these data indicate that Creb3l2 is acting very broadly on a large number of translation regulatory and effector genes in order to enhance translation capacity. This situates the Creb3l2 cell-autonomous action downstream of the signaling pathways that are usually implicated in translational control, such as the Akt/mTOR pathway.

### Creb3l2, a translation scaling factor

To evaluate to which extent gene expression changes dependent on Creb3l2 or/and XBP1 resemble gene expression changes caused by *Tpit* inactivation, we performed RNAseq on our LOF and GOF models and analyzed the results by unsupervised clustering analysis. Four gene clusters (A-D) were identified and analyzed by Gene Ontology (GO) (Fig. 5a). Two clusters appeared to be particularly interesting: clusters A and C. Cluster A is enriched in genes controlling protein transport and translation that tend to be downregulated in *Tpit* KO tissues. The same tendency is observed for these genes in ACreb3l2 and AXBP1 transgenic mice, and even more in combined ACreb3l2/AXBP1 mice. Cluster A includes about 50% of the genes in each translation sub-category (Fig. 4g). Hence, combined downregulation of Creb3l2 and XBP1 phenocopies the effect of Tpit deficiency on translation and secretory pathway genes.

Cluster C represents a group of genes synergistically downregulated in Creb3l2+XBP1 cells and it is enriched in genes controlling hexose metabolism. Their analysis suggested a shift from glycolysis to oxidative phosphorylation utilization of glucose and in agreement with this, mRNA levels of lactate dehydrogenase (Ldha) and pyruvate dehydrogenase kinase (Pdk1) are decreased in Creb3l2+XBP1 cells (Fig. 5b). Accordingly, the latter cells exhibit decreased lactate production and glucose utilization (Fig. 5c). We directly assessed cell respiration and found increased ATP turnover only in Creb3l2+XBP1 cells (Fig. 5c). Since AtT-20 cells are tumor-derived, they are likely to exhibit a Warburg shift towards glycolytic glucose utilization compared to normal pituitary cells. The Creb3l2-dependent reverse shift towards the more energy efficient mitochondrial oxidative phosphorylation pathway is consistent with the high energy requirements for translation and the development of a high capacity secretory system. In order to assess conservation of Creb3l2 and UPR regulators in an evolutionary distant but related system, we queried their status in the *Xenopus* IL where melanotrope cell size and αMSH secretion increase when frogs adapt to a dark background environment ^17,32^. We found that IL RNA content (Fig. 5d) and expression levels of UPR pathway regulators, including Creb3l2, are increased in dark-adapted frogs (Fig. 5e). Thus, the physiological UPR pathway appears an evolutionary conserved mechanism to match secretory capacity with translational burden.

**Figure 5.**
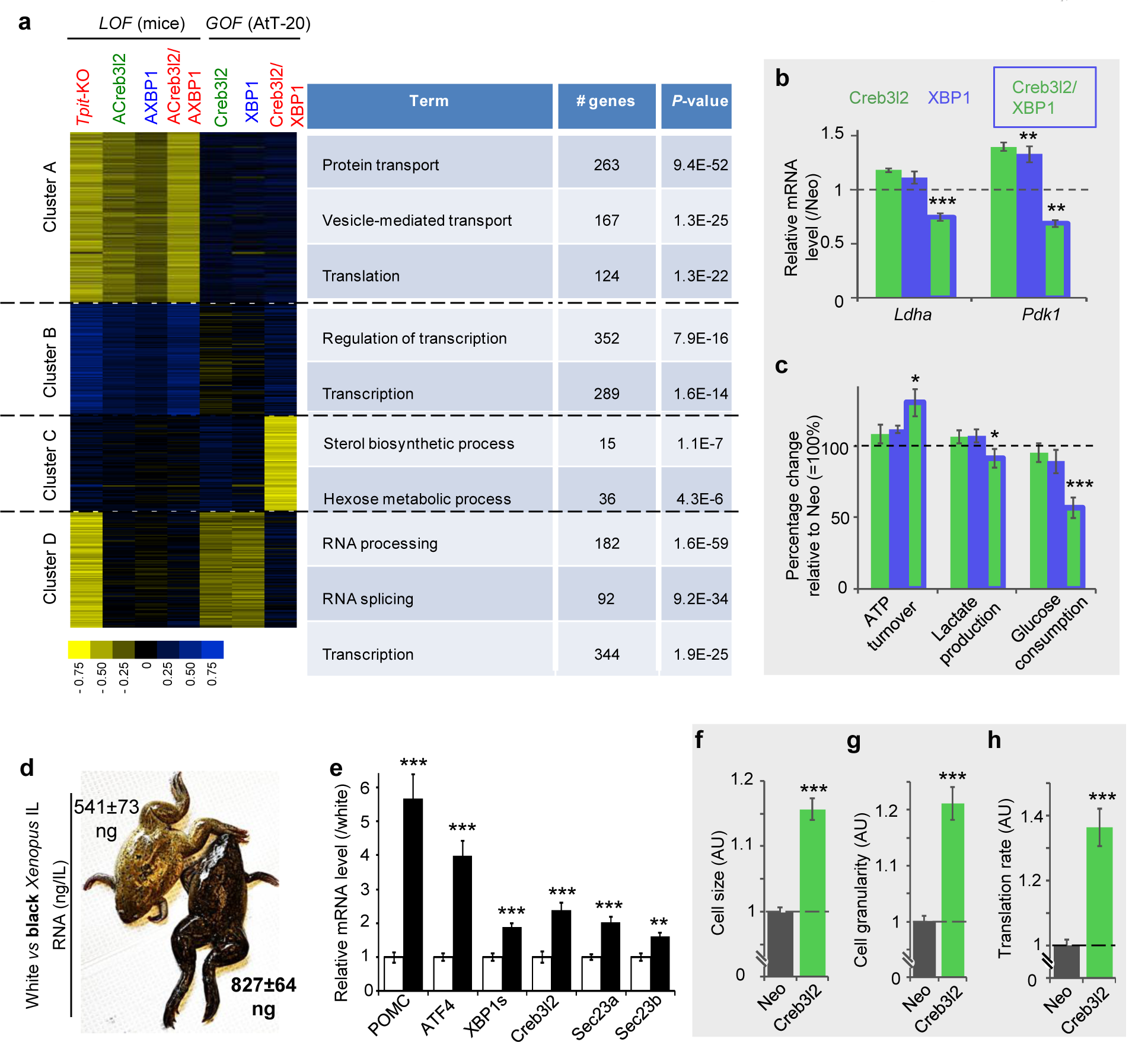
Creb3l2 is a scaling factor for translation. (**a**) *Tpit* KO transcriptome signature is largely phenocopied by inhibition of *Creb3l2* and *XBP1*. Heatmap representation of gene clustering identified by global analysis of expression profiling datasets (*left*) and GO distribution of genes in each cluster (*right*). Transcriptome data from WT, *Tpit-KO*, *ACreb3l2*, *AXBP1* and *ACreb3l2/AXBP1* transgenic mice together the gain-of-function AtT-20 cells were clustered with the Cluster 3.0 software using K-means. (**b**) Decreased lactate dehydrogenase (Ldha) and pyruvate dehydrogenase kinase (Pdk1) mRNAs (RNAseq normalized counts) in AtT-20 cells expressing Creb3l2+XBP1 relative to control (Neo) cells set to 1. (**c**) Increased ATP turnover, decreased lactate production and glucose utilization, showing shift of ATP production from glycolysis to oxidative phosphorylation in AtT-20 cells expressing Creb3l2+XBP1. Data represent the means ±SEM (n= 5-6). (**d**) Adaptation (5 weeks) of *Xenopus* skin color to black (*right*) background compared to white (*left*) increases total IL RNA content (ng ±SEM, for n= 4-5). (**e**) Expression of *Pomc* and UPR genes in white (white bars) and black (black bars) background adapted frogs. Relative IL mRNA levels measured by RT-qPCR were normalized to those of β*-actin* and represented as fraction of levels in white background set to 1. Data represent means ±SEM (n= 4-5). (**f-h**) Expression of Creb3l2 in INS-1 cells. FACS analyses of cell size (**f**) and granularity (**g**) index for INS-1 cells expressing Creb3l2. All data represent the means ±SEM (n = 3). (**h**) SUnSET measurement of protein synthesis in control INS-1 cells (Neo) and INS-1 cells expressing Creb3l2. Data (±SEM, n = 3) are presented relative to Neo cell set to 1.

To our knowledge, Creb3l2 is the first TF shown to directly target and regulate the translation transcriptome. Creb3l2 is expressed in many secretory tissues (Supplementary Fig. S1f) including in pancreatic islets. While pituitary POMC cell maturation is regulated by Tpit, pancreatic β cell maturation is dependent on the TF Pdx1. Interestingly, Pdx1 binding is observed at the *Creb3l2* promoter in islet cells (Supplementary Fig. S4e) suggesting a maturation mechanism that may resemble that of POMC cells. To assess Creb3l2 activity in β cells, we stably overexpressed Creb3l2 in insulin-secreting INS-1 cells and found an increase of INS-1 cell size, granularity and a 36% increase of translational rate (Fig. 5f-h). Thus, Creb3l2 appears to be a general regulator of translation capacity.

## Discussion

The present work identified a master TF for control of translation capacity, Creb3l2. Genome-wide approaches demonstrated that Creb3l2 is directly targeting and stimulating transcription of hundreds of genes involved in different steps of translational control, namely 55 translation factors, 105 ribosomal proteins and 30 aminoacyl tRNA synthetases genes. Genetic manipulations (LOF, GOF) of the *Creb3l2* gene in mice and two different secretory cell types led to significant changes of translation rates (25-36%). Overexpression of Creb3l2 was sufficient to increase overall cellular protein content by ≈25%. Scaling factors were proposed to be factors that adjust expression of genes controlling basic cell functions to specific cellular needs ^33^. Creb3l2 is thus a scaling factor for translation.

It is noteworthy that total RNA content is increased following Creb3l2 action. Although we could not find evidence of direct Creb3l2 binding to rRNA genes, its broad action on expression of ribosomal proteins is likely sufficient to increase rRNA transcription and hence, have a significant effect on total RNA content. It was indeed shown that depletion of ribosomal proteins RPS19, RPS6 or RPL11 is sufficient to decrease rRNA transcription ^34^.

The present work suggests a general paradigm that involves three TFs for maturation of secretory cells into hormone-producing factories (Fig. 6). A tissue-specific terminal differentiation factor (Tpit in pituitary POMC cells, Pdx1 in pancreatic islet cells) that upregulates two bZIP TFs, Creb3l2 for scaling up translation capacity and XBP1 to increase secretory organellogenesis without triggering the stress response. These physiological regulators of secretory capacity could be useful to increase yields of therapeutic proteins such as cytokines and monoclonal antibodies that are produced by expression in mammalian cells.

**Figure 6.**
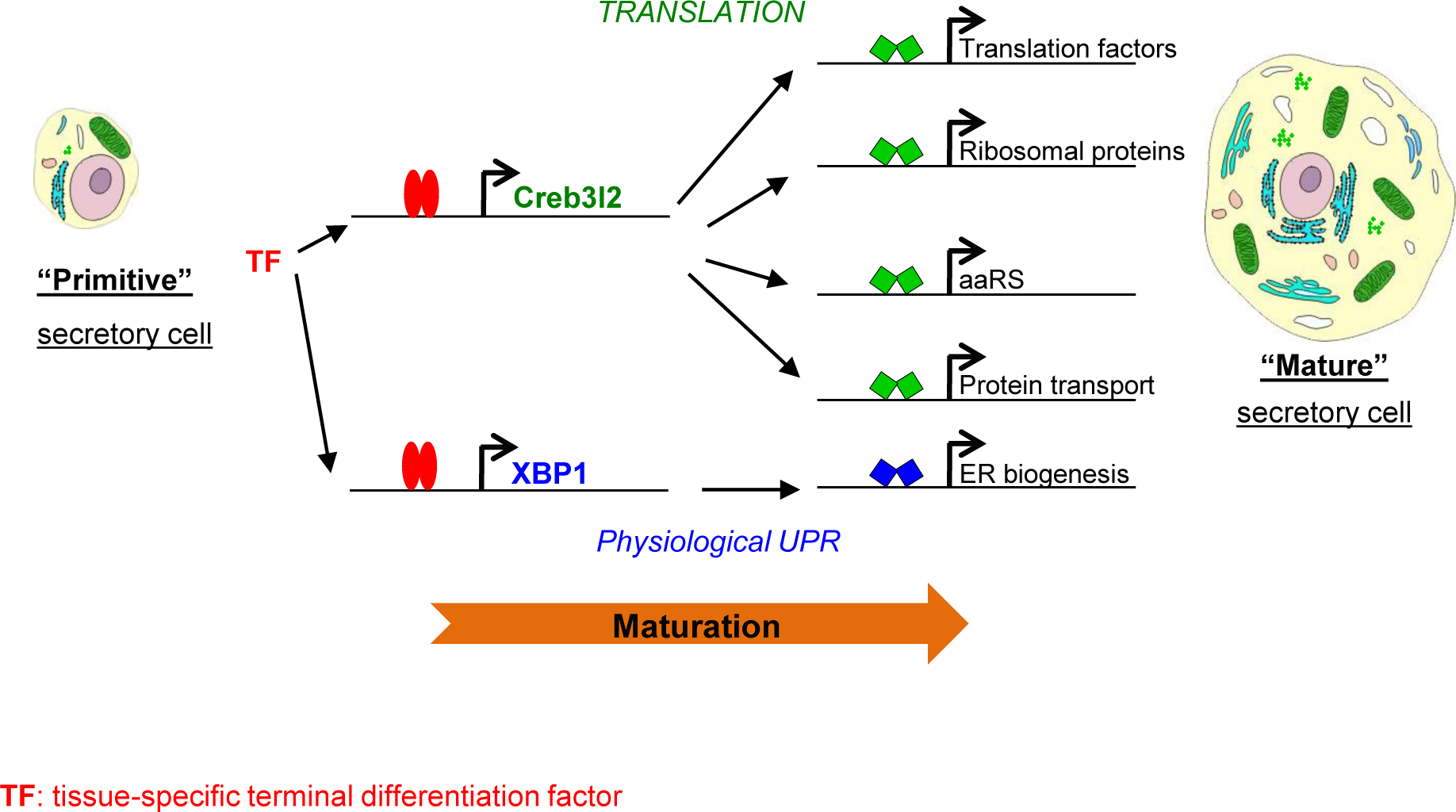
Mechanism for setting high secretory capacity. Two bZIP transcription factors Creb3l2 and XBP1 transform differentiated secretory cells into hormone-producing factories by scaling up translation (Creb3l2) and activating the physiological UPR pathway (XBP1) to prevent protein stress response. The expression of both Creb3l2 and XBP1 is increased by tissue-specific differentiation TFs, Tpit in pituitary POMC cells and Pdx1 in pancreatic β cells.

The involvement of the UPR regulator XBP1 in this regulatory network clearly fits within the context of the so-called “physiological”UPR” system rather than within the cytoprotective homeostatic response to protein stress. The “physiological UPR” was described in professional secretory cells, such as B lymphocytes, digestive enzyme-secreting zymogenic cells, salivary glands, exocrine pancreas or pancreatic beta cells ^1,35–37^. This proactive UPR engagement is preparing mature secretory cells to massive protein synthesis and secretion, rather than being a response to high secretory load. The existence of two types of UPR (i.e. stress related and physiological) raises the question of how context-dependent UPR specificity is achieved. The stress-related UPR involves three sub-pathways characterized by different ER transmembrane sensor proteins, IRE1, PERK and ATF6. A clearly undesirable aspect of UPR in highly secreting cells is the PERK branch that leads to global translational repression by phosphorylation and inhibition of eIF2α ^38^. Interestingly, PERK expression is barely affected by loss of Tpit but expression of Gadd34/Myd116, a phosphatase reversing eIF2α phosphorylation is positively regulated by Tpit. In fact, Tpit action not only seems to exclude PERK, but also prepare POMC cells to neutralize PERK activation. Also, genes related with ER associated protein degradation (ERAD genes) are mostly unaffected in *Tpit-/-* pituitaries. Another undesirable UPR effect in physiological conditions is the apoptotic response that follows prolonged ER stress (reviewed in ^39^). In *Tpit-/-* IL, the expression of CHOP/Ddit3, a major mediator of ER stress-induced apoptosis is not affected. During ER stress, apoptosis can be delayed through expression of inhibitors of apoptosis (IAPs) proteins ^40–42^. Expression of two IAPs (Naip5 and Naip6) is strongly dependent on Tpit and Tpit binding is observed at these genes (not shown). Thus, Tpit protects POMC cells from apoptotic signals. Collectively, these results suggest that Tpit sets up an efficient secretory system by activating “useful” (XBP1, Creb3l2) but not detrimental (PERK, CHOP/Ddit3) UPR regulators and stimulating inhibitors of deleterious UPR effects (the eIF2α phosphatase Gadd34/Myd116 and inhibitors of apoptosis Naip5 and Naip6).

Importantly, in addition to XBP1 stimulation of ER biogenesis, XBP1 has a synergistic effect on protein synthesis in combination with Creb3l2, increasing the overall cellular protein content by ≈50% with resulting enhancement of regulated secretion. Translation is the most energy consuming (about 70%) cellular process ^43^ and uncontrolled protein synthesis leads to apoptosis ^44^; in the latter work on translational activation through the PERK pathway mediated by ATF4 and CHOP ^44^, it is noteworthy that the group of stimulated translation control genes included primarily tRNA aminoacylation in contrast to the broad set identified in the present work. Remarkably, the combined Creb3l2 plus XBP1 expression in the tumor-derived AtT-20 cells led to a shift of energy metabolism from glycolysis to oxidative phosphorylation, resulting in more efficient ATP production and providing for support of heavy protein synthesis.

In summary, the present work identified Creb3l2 as a scaling factor for translation and showed that its joint action with XBP1 dramatically increases protein synthesis and secretory capacity. The targets of Creb3l2 and XBP1 action are revealing of their purpose. By targeting genes that are downstream within the Creb3l2 regulatory network (Fig. 6), this TF sets the cell-autonomous “rheostat” for translation capacity, in this instance setting high level capacity. And the joint action of XBP1 on the physiological UPR pathway matches this with increased secretory capacity. This reset of protein secretion capacity is separate from, and therefore compatible with, upstream signaling pathways regulating translation such as the Akt/mTOR pathway.

## Acknowledgements

K.K. dedicates this manuscript to the memory of Régine Losson who died untimely 8 years ago. She was a wonderful person, an outstanding scientist and a great mentor. We are grateful to our colleagues Nahum Sonenberg, John J Bergeron, Nicole Francis and Steve Bilodeau for comments on the manuscript. We thank David Langlais, Lionel Budry and Benjamin Haibe-Kains for helpful discussions, Eric Massicotte for help with FACS, Dominic Filion for the script allowing image analysis, Amandine Bemmo, Alexis Blanchet-Cohen and Nicolas Bouchard for the help in RNAseq analyses. This work was supported by grants from Canadian Institutes of Health Research (CIHR MOP-86650 and FRN-154297) and by access to Compute Canada resources. KK was supported by CIHR and FRQS.

## Author contributions

K.K. performed experiments. K.K. and J.D. conceived and designed the experiments. A.B. and A.M. helped with experiments and ChIPseq data analysis. H.C. performed and analyzed EM. J.SP. and V.C. conceived and performed respiration tests. K.K. and J.D. wrote the manuscript.

## Methods

### Animals and genotyping

The generation and genotyping of *Tpit* KO (BALB/c) and POMC-EGFP transgenic (C57BL/6) mice was described elsewhere ^15,20^. *Pomc*-KO (C57BL/6) mice ^45^ were purchased from Jackson laboratory (B6.129X1-Pomctm2Ute/J strain, stock number 008115). Young-adult (6 months-old) *Xenopus laevis* were kindly provided by Dr Marko Horb. Frogs were fully adapted to black or white background (6 animals per condition) for 5 weeks. After adaptation, animals were anesthetized by submersion in a solution of benzocaine (0.05%), rapidly decapitated, and their IL dissected, frozen in liquid nitrogen and kept at −80°C until RNA extraction.

#### Transgenesis

DNA fragments containing ACreb3l2 or AXBP1 dominant-negative cDNA constructs under the control of rat *Pomc* promoter and followed by a simian virus 40 (SV40) intron and polyadenylation sequences were microinjected and founder lines were derived as previously described ^15^.

### Histology and electron microscopy

Processing of PFA-fixed, paraffin-embedded pituitaries and immunohistofluorescence were done as described ^46^. For electron microscopy, adult (4-6 month-old) mice were first perfused with PBS without potassium (10 ml/mice at 2 ml/min), then with 2.5% glutaraldehyde (20 ml/mice at 2 ml/min). Pituitaries were then dissected and post-fixed in 2.5% glutaraldehyde for 2h at +4°C. EM was performed as previously described ^47^. Briefly, the tissue was contrasted with uranyl acetate [2% (w/v) in distilled water], dehydrated in ethanol and embedded in LR Gold resin (Agar Scientific, London, UK). Ultrathin sections (50-80 nm) were prepared using a Reichart-Jung ultracut microtome, mounted on nickel grids (Agar Scientific, Stanstead, Essex, UK) and examined on a JOEL 1010 transmission electron microscope (JOEL USA Inc., Peabody, MA, USA).

#### Quantitative Electron Microscopy

For analysis of cell morphology ten micrographs of intermediate lobe cells per animal (n=4 mice per group) were taken at a magnification of x 4,000 and scanned into Adobe Photoshop (version 5.5) and analyzed using Axiovision (version 4.5) image analysis software. The analyst was blind to the sample code. The following parameters were calculated: cytoplasmic, nuclear and total cell areas; granule area, granule density, and granule diameter. For measurement of the cell and nuclear areas, margins were drawn around the cell or nucleus respectively and the area was calculated. Cytoplasmic area was determined by substracting nuclear area from total cell area. Granule density was calculated by dividing total granule area by cytoplasmic area. Expansion of the rough endoplasmic reticulum (RER) and Golgi apparatus was assessed visually and graded on a scale of 0-4 (0, no expansion; 4 the most expansion). These estimates do not provide absolute measurements but do provide a basis for comparison.

### IL cell (melanotrope) analyses

#### DNA content

IL were first digested in 1.5 ml tubes with proteinase K (PK: 30 µg) in 400 µl of PK-lysis buffer (0.5% SDS - 100 mM NaCl, 50 mM Tris, pH 7.5 - 1 mM EDTA) at 55°C, overnight. After a brief RNase treatment (15 min), ILs were redigested with PK (20 µg, 55°C, 1 h). Genomic DNA was then precipitated with 1 volume of 4 M ammonium acetate – 0.6 volume isopropanol, washed with 70% ethanol and dissolved in 20 µl of 10 mM Tris, pH 8.0 at 50°C, overnight. DNA was quantitated using TKO100 Fluorometer, following manufacturer’s operating instructions (Hoefer Scientific Instruments) in order to specifically quantify double stranded nucleic acids.

#### RNA content

Total RNA from pools of 5 ILs was extracted using RNeasy Plus Mini kit (Qiagen, 74134) following manufacturer’s recommendations, including the step of genomic DNA removal. Experiments were repeated on pools of 5-9 ILs at least 3 times. Statistical significance was assessed using bilateral Student’s *T*-test with unequal variances on Microsoft Excel.

#### Translation rate

Translation monitoring was done using the principles of surface sensing of translation (SUnSET) ^27^ adapted for *in vivo* studies ^48^. Briefly, adult (3-6 months) mice were intraperitoneally injected with 100 µl of 7 mg/ml puromycin dihydrochloride (Gibco A1113803) solution (0.04 µmol/kg). 30 min after injection ILs were extracted, and melanotrope cells prepared as described ^49^. After fixation and permeabilization (BD Cytofix/Cytoperm 554714) cells were stained in 96-well plates with anti-Puromycin Alexa647 antibodies (mouse monoclonal, Millipore MABE343-AF647, dilution1/75) in 30 µl of 1xPBS – 0.5% BSA (+4°C, 30 min). Cells were then washed in 1xPBS – 0.1% BSA, resuspended in washing buffer and analyzed by FACS. The experiment was repeated on pools of 4-6 ILs at least 5 times. Statistical significance was assessed using bilateral Student’s *T*-test with unequal variances on Microsoft Excel.

#### ER content

Melanotrope cells were prepared as described above. Living cells were then stained with 1 µM ER-tracker green (ThermoFisher Scientific E34251) in 30 µl HBSS (96-well plates) at 37°C, 5% CO_2_. Cells were then washed in 1xPBS – 0.1% BSA, resuspended in the same washing buffer and analyzed by FACS. The experiment was repeated on pools of 3-5 IL at least 3 times. Statistical significance was assessed using bilateral Student’s *T*-test with unequal variances on Microsoft Excel.

#### Nuclear density

Pituitary sections were stained with anti-Tpit antibodies (homemade rabbit polyclonal 1250B, dilution 1/100) or with Hoechst (SIGMA B2883) in the case of *Tpit*−/− pituitaries. After image acquisition IL nuclear density was analyzed using the Imaris Software and custom written MATLAB programs designed as follows: (1) the region was first outlined by hand to keep only the pituitary, (2) the position of each cell was identified via the fluorescent channel using the spot function from Imaris with background subtraction and fixed quality factor, (3) the area was calculated from the object thus created and, (4) cell density was counted as the number of spots over the object surface. The full processing *XTension* method is found at http://open.bitplane.com/Default.aspx?tabid=57&userid=337 with an example. Statistical significance was assessed using bilateral Student’s *T*-test with unequal variances on Microsoft Excel.

### Cell culture

#### Pituitary cell lines (mouse)

AtT-20, αT3 or GH3 cells were cultured in Dulbecco’s modified Eagle’s medium supplemented with 10% fetal bovine serum and antibiotics (penicillin/streptomycin).

#### Pancreatic insulinoma cells (rat)

INS-1 cells were cultured in 1x RPMI1640 medium (Wisent 350000CL) supplemented with 10 mM HEPES, pH 7.4, 1 mM Na pyruvate, 50 µM β-mercaptoethanol, 10% fetal bovine serum and antibiotics (penicillin/streptomycin).

#### Generation of stable transgenic cell populations

Retroviruses were packed using the EcoPack 2-293 cells (Clontech, Mountain View, CA) and infections were performed as described ^50^. Selection of retrovirus-infected cell populations was achieved with either 400 µg/ml Geneticin (Gibco, 11811-031) or 200 µg/ml Hygromycin B (Invitrogen, 10687-010). Resistant colonies were pooled to generate retrovirus-infected populations of about 1,000 independent colonies. For double infections, cells were first infected by pLNCX2/3xFLAG-Creb3l2 and the resulting “Creb3l2/Neo” cells were re-infected by pLHCX/XBP1s and selected on media containing Geneticin/Hygromycin.

For genomic DNA extraction and quantification, AtT-20 cells were seeded in triplicates in 12-well plates (3×10^5^ cells/well) at day 0. At day 2, media were changed and at day 3, genomic DNA was extracted as for IL samples, resuspended in 50 µl of 10 mM Tris, pH 8.0 and quantitated using a fluorometer.

Total RNA was extracted using RNeasy Mini kit (Qiagen, 74104) following manufacturer’s recommendations. Experiments were done in triplicates. Statistical significance was assessed using bilateral Student’s *T*-test with unequal variances on Microsoft Excel.

For measurements of protein content, cells were plated as for DNA quantification. At day 3, cells were washed twice with 1xPBS, lysed with the EBC buffer (50 mM Tris-HCl pH 8.0, 170 mM NaCl, 0.5% NP-40, 50 mM NaF, 10% glycerol) and soluble protein concentration was measured by absorbance using the Bradford assay (Bio-Rad protein assay 500-0006). For each cell line, 4 wells were used for protein extractions and 3 for genomic DNA quantitation (as described above), used as indicator of cell number. The experiment was performed in quadruplicates 5-8 times. Statistical significance was assessed using bilateral Student’s *T*-test with unequal variances on Microsoft Excel.

ACTH release was measured using a commercially available ELISA kit (**mdb**ioproducts M046006). AtT-20 cells grown on 12-well plates (3×10^5^ cells/well) for 48h were placed on lipid-free low-serum medium (0.5% Dextran-coated charcoal treated FBS) for 12h. Cells were then washed with serum-free medium and placed in 1 ml of serum-free medium containing CRH (10^-7^ M) or vehicle (1xPBS) for 7h. The accumulated amount of ACTH released in the medium was measured following instructions of the manufacturer using 0.4 µl of medium (100 µl of a 1/250 dilution) per assay. Genomic DNA (quantified as above) was used as indicator of cell number. The experiment was done in duplicates at least 3 times. Statistical significance was assessed using bilateral Student’s *T*-test with unequal variances on Microsoft Excel.

Metabolic studies in different AtT-20 cell populations (respiration, lactate and glucose levels) were performed as described ^51^.

#### Transient transfections

αT3 (Fig. 2g), GH3 (Fig. 4f and Suppl. Fig. S4c,d) or INS-1 (Fig. 3b and Suppl. Fig. S2a,b) cells were plated in 12-well plates (3×10^5^ cells/well) the day preceding transfection and transfected with 1.4 µg of DNA, containing 100 ng of luciferase reporter construct with different amounts of expression vectors completed with empty expression vector (JA1394) using Lipofectamine reagent (Invitrogen, Carlsbad, CA).

### DNA Vectors

Supplementary Table S3 lists reagents used to construct all expression vectors, including relevant PCR primers and used restriction sites.

#### Retroviral vectors (for generation of stable cell populations)

The following retroviral vectors were used: pLNCX2 (JA1784), pLNCX2/XBP1s (JA2335), pLHCX (Clontech), pLHCX/XBP1s (JA2340) and pLNCX2/3xFLAG-Creb3l2 (JA2349).

#### Transient transfection vectors

XBP1s and Creb3l2 (cleaved fragment) expression vectors (JA2343 and JA2426, respectively) were constructed in the same expression vector (JA1394) as previously described for Tpit ^18^, containing the T3 promoter, allowing production of radiolabelled proteins *in vitro* (see pull-down description). MBP-βGal (pMAL-C) and MBP-Tpit (JA1250) expression vectors were previously described ^18^. The coding sequences corresponding to XBP1s (*spliced* XBP1) and Creb3l2_CL_ (*cleaved* Creb3l2) were amplified from total AtT-20 cDNA by PCR. To obtain the 3xCreb3l2-RE-luciferase reporter (JA2422), we hybridized oligonucleotides 59-5/59-6 and then cloned their double stranded DNA into the BamHI site located upstream of a minimal POMC promoter in plasmid JA290 ^52^.

#### Dominant-negative (DN) expression vectors (for transgenesis in vivo)

First, the amphipathic acidic extension (AZIP) from the previously described CMV A-CREB construct (Addgene catalog #33371) ^53^ was PCR amplified with primers 30-585/25-208, introducing a NotI restriction site on 5’ end and an XhoI site at the 3’ end. Second, Creb3l2 and XBP1 coding sequences starting at the leucine zipper domain were PCR amplified with primers 31-245/36-147 and 32-287/33-263 introducing a XhoI restriction site on 5’ end and an NotI site at the 3’ end. The, NotI-AZIP-XhoI DNA fragment was ligated to XhoI-Creb3l2-NotI or XhoI-XBP1-NotI and the resulting NotI-ACreb3l2-NotI or NotI-AXBP1-NotI fragments were subcloned in the vector JA1394/NotI, giving rise to plasmids JA2428 and JA2429, respectively. Finally, NotI-ACreb3l2-NotI or NotI-AXBP1-NotI DNA fragments from JA2428 and JA2429 were blunt-ended with DNA polymerase I large fragment (Klenow) and cloned into JA1508/EcoRI-Klenow downstream of the rat *Pomc* promoter and upstream of the SV40 intron. The resulting plasmids JA2452 and JA2453 were digested by SalI to generate SalI-POMC_promoter-ACreb3l2-SV40-SalI (≈1.8 kb) and SalI-POMC_promoter-AXBP1-SV40-SalI (≈2.5 kb) DNA fragments used for oocyte microinjection.

### FACS analysis

IL cells were prepared and analyzed using the FACS Calibur cell sorter (BD BioSciences) as described ^49^. Propidium iodide labeling (1 µg/10^6^ cells) was used to exclude dead cells. Data were analyzed with the Summit 4.3 software. Cell diameter of IL cells was inferred by extrapolating forward scatter fluorescence values to the standard curve made using a set of calibrated beads (Spherotech PPS-6K). Cell granularity was assessed based on the side scatter (SSC) measured by FACS. Size and granularity of AtT-20 and INS-1 cells were analyzed the same way. The experiments were repeated at least 3 times. Statistical significance was assessed using bilateral Student’s *T*-test with unequal variances on Microsoft Excel.

### RNA extraction, cDNA synthesis and transcriptomic analyses

IL or AtT-20 cell RNA was extracted using RNeasy mini extraction columns (Qiagen) according to the manufacturer’s instructions. 100-200 ng (IL) of total RNA were utilized to generate cDNA using the Superscript III reverse transcriptase (Invitrogen, 18080-044) following the manufacturer’s recommendations. Resulting cDNAs were analyzed by qPCR using a SYBR Green master mix (ThermoFisher Scientific A25741) supplemented with 300 nM of each gene specific primer pair (see Suplementary Table S4 for sequences). Transcriptomic (RNAseq) studies were performed in triplicates for different AtT-20 cell populations or in duplicates for mice, using pools of five 3 months-old male IL tissues. For *WT* or *Tpit-KO* ILs, cDNA probes were hybridized on Affymetrix mouse Gene 1.0 ST arrays. Hybridization and scanning were done at the McGill University and Genome Québec Innovation Centre. RNASeq were done at the IRCM Molecular Biology facility following the TruSeq Stranded mRNA sample preparation guide (Illumina).

### Chromatin Immunoprecipitation – sequencing (ChIPseq)

ChIP experiments were performed in AtT-20 cells as previously described ^54^ using 5 µg of antibody (anti-FLAG-M2, mouse monoclonal, Sigma F3165; anti-mouse XBP1, rabbit polyclonal M-186, Santa Cruz Sc-7160) per reaction. Equal amounts of purified rabbit or mouse IgG (rabbit IgG, Sigma G2018; mouse IgG, Sigma I5381) were used in control ChIP reactions. Pools of DNA (80-100 ng) from several independent ChIP experiments were utilized to make libraries for ChIPseq. The libraries and flowcells were prepared at the IRCM Molecular Biology Core facility following Illumina recommendations (Illumina, San Diego, CA) and then sequenced on Hiseq 2000 at the Génome Québec Innovation Centre. The generation of Tpit and H3K4me1 ChIPseq and FAIREseq profiles in AtT-20 cells was already described ^22^. H3K27ac ChIPseq is described ^55^ and available on GEO as GSE 87185.

### Bioinformatics processing and analysis of genomic data

The FlexArray software developed at McGill University (http://genomequebec.mcgill.ca/FlexArray) was used to analyze transcriptomic data (GC-RMA normalization and Scatter plots). Differentially expressed genes were extracted by fixing a *P*-value threshold (*P*≤0.001) using the Local-pooled-error (LPE) test.

Finding of enriched functional-related gene groups was done using Gene Ontology tool from the Database for Annotation, Visualization and Integrated Discovery (DAVID) ^56^ or AmiGO1 (http://amigo1.geneontology.org/cgi-bin/amigo/term_enrichment) ^57,58^. *P*-values on DAVID (Fig. 2b) correspond to EASE Score, a modified (more conservative) Fisher Exact *P*-value, while *P*-values on AmiGO (Fig. 4e) were calculated with GO::TermFinder using the hypergeometric distribution. The heat map of transcriptomic microarray data was visualized using the TreeView Java application ^59^.

ChIPseq peak finding and sequence analysis were described previously ^22^. For *de novo* motif analysis, 120 bp of sequences surrounding the peaks were extracted from the University of California at Santa Cruz Web site ^60^ and processed using HOMER ^61^. It uses ZOOPS scoring (zero or one occurrence per sequence) coupled with the hypergeometric enrichment calculations (or binomial) to determine motif enrichment. Graphical representations of the position weight matrices obtained from these analyses were generated with WebLogo (http://demo.tinyray.com/weblogo) ^62^. Peaks were assigned to the closest gene with the AnnotatePeaks.pl Homer command. To visualize ChIPseq profiles, we used the Integrative Genome Viewer tool ^63^.

RNAseq reads were trimmed with Trimmomatic v0.22 ^64^ and aligned against the mm10 mouse genome using Tophat (v2.0.8). Raw read counts per gene were calculated with HTSeq-count from HTSeq (v0.5.4) (http://www-huber.embl.de/users/anders/HTSeq/). The raw counts were normalized (N) relative to the library size, and tested for differential expression using the R package DESeq ^65^. Genes were considered to be expressed if the average normalized count across the samples was above 20.

#### Unsupervised clustering

A table containing gene expression fold changes in different LOF or GOF conditions was uploaded into Cluster 3.0 software (enhanced version of Cluster, originally described in ^66^). After normalization, we applied K-means for clustering using Euclidean distance similarity metrics. We determined that 4 clusters is the most segregating setting for our dataset. The gene lists extracted from those 4 clusters were uploaded into the DAVID website ^56^ to search for enriched biological processes.

## Supplemental Figure Legends

**Figure S1.**
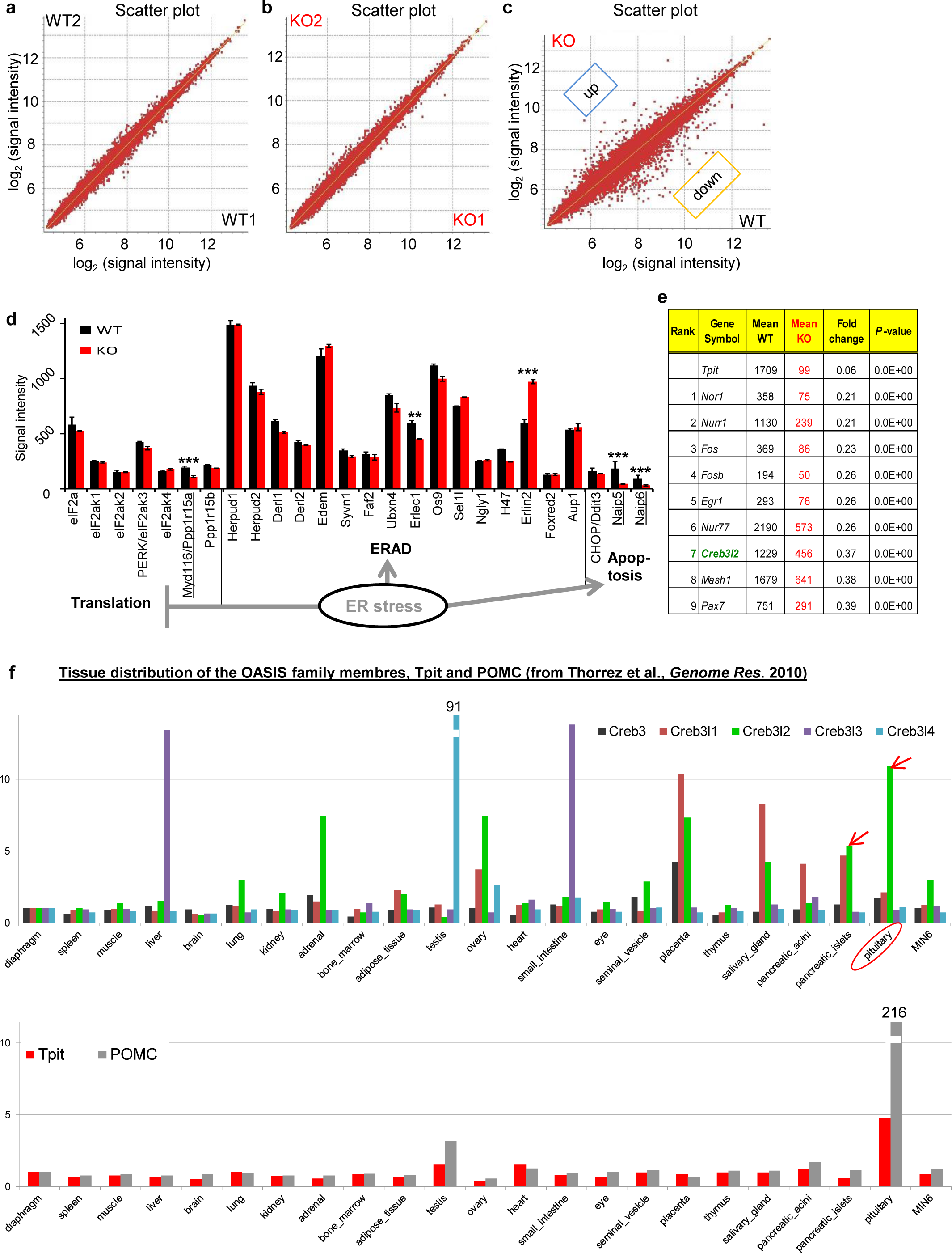
Validation of Tpit KO (Tpit-/-) transcriptome data. (**a-c**) Scatter plot comparisons of 2 WT (**a**) and 2 KO (**b**) pituitary intermediate lobe (IL) transcriptomes showing the reproducibility of microarray experiments and comparison of WT *vs* Tpit KO pituitary IL transcriptomes (**c**). (**d**) Expression (Affymetrix microarray signals) of translation regulators, ERAD and apoptotic genes in WT (black bars) and KO (red bars) ILs. (**e**) List of 10 most downregulated transcription factor genes in *Tpit*-deficient ILs. (**f**) Expression of the OASIS family members, *Tpit* and *POMC* in different tissues and cell lines.

**Figure S2.**
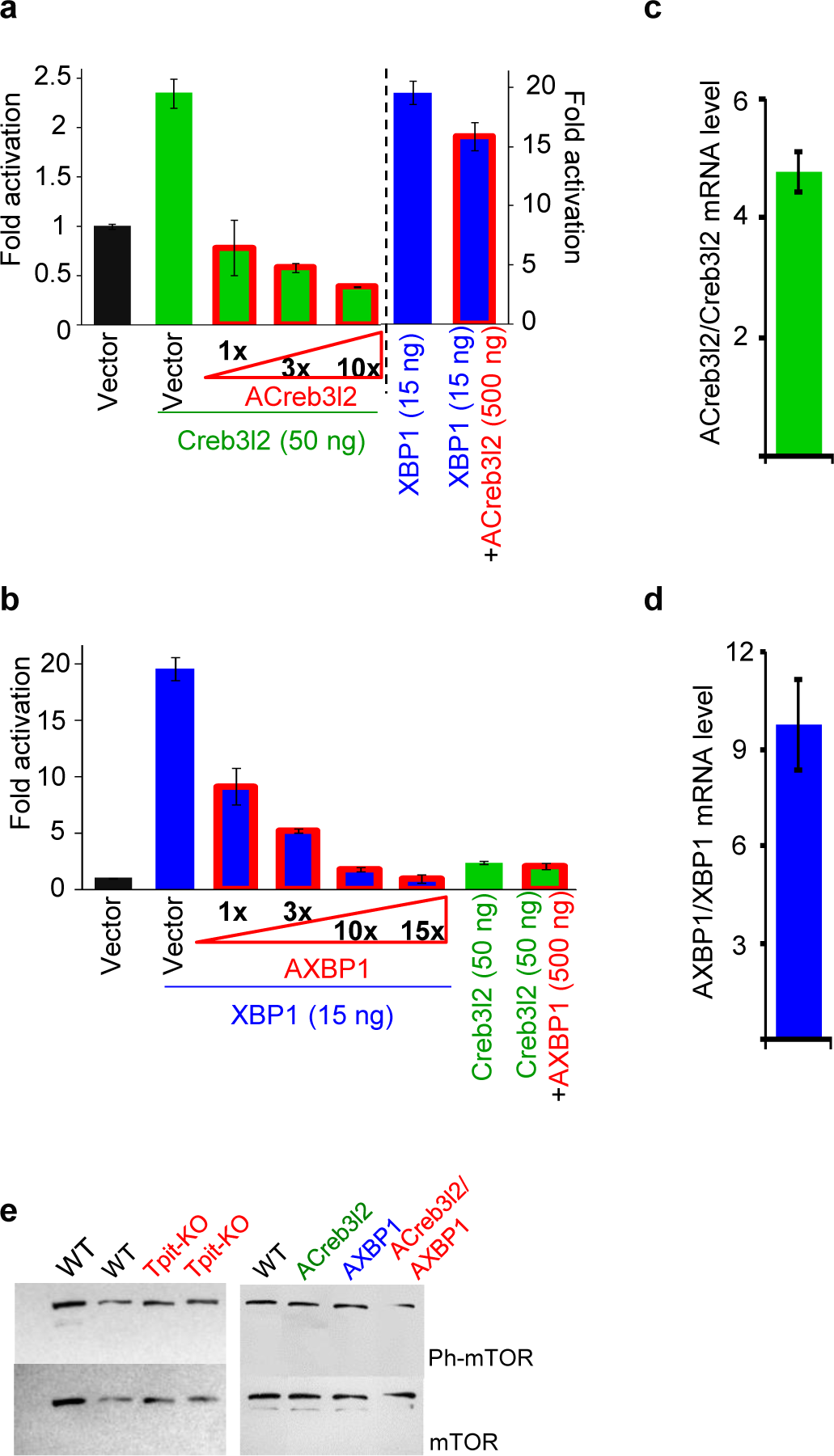
Efficient and specific inhibition of Creb3l2 and XBP1 activity using acidic dominant-negative constructs (AZIP). (**a**) Inhibition of Creb3l2, but not XBP1, activity by overexpression of ACreb3l2. (**b**) Inhibition of XBP1, but not Creb3l2, activity by overexpression of AXBP1. The 3xCreb3l2/XBP1-RE *Luciferase* reporter was used to assess Creb3l2 or XBP1 activity upon transfection into INS-1 cells. (**c-d**) RT-qPCR quantification of ACreb3l2 and AXBP1 transcripts relative to the endogenous Creb3l2 and XBP1s transcripts in ILs of the transgenic mice reported in Fig. 3. (**e**) mTORC1 status is not affected by Tpit KO or Creb3l2/XBP1 downregulation (ACreb3l2, AXBP1). Western blot showing similar active mTOR (phospho Ser-2481) to total mTOR levels in IL protein extracts from indicated mutant mice.

**Figure S3.**
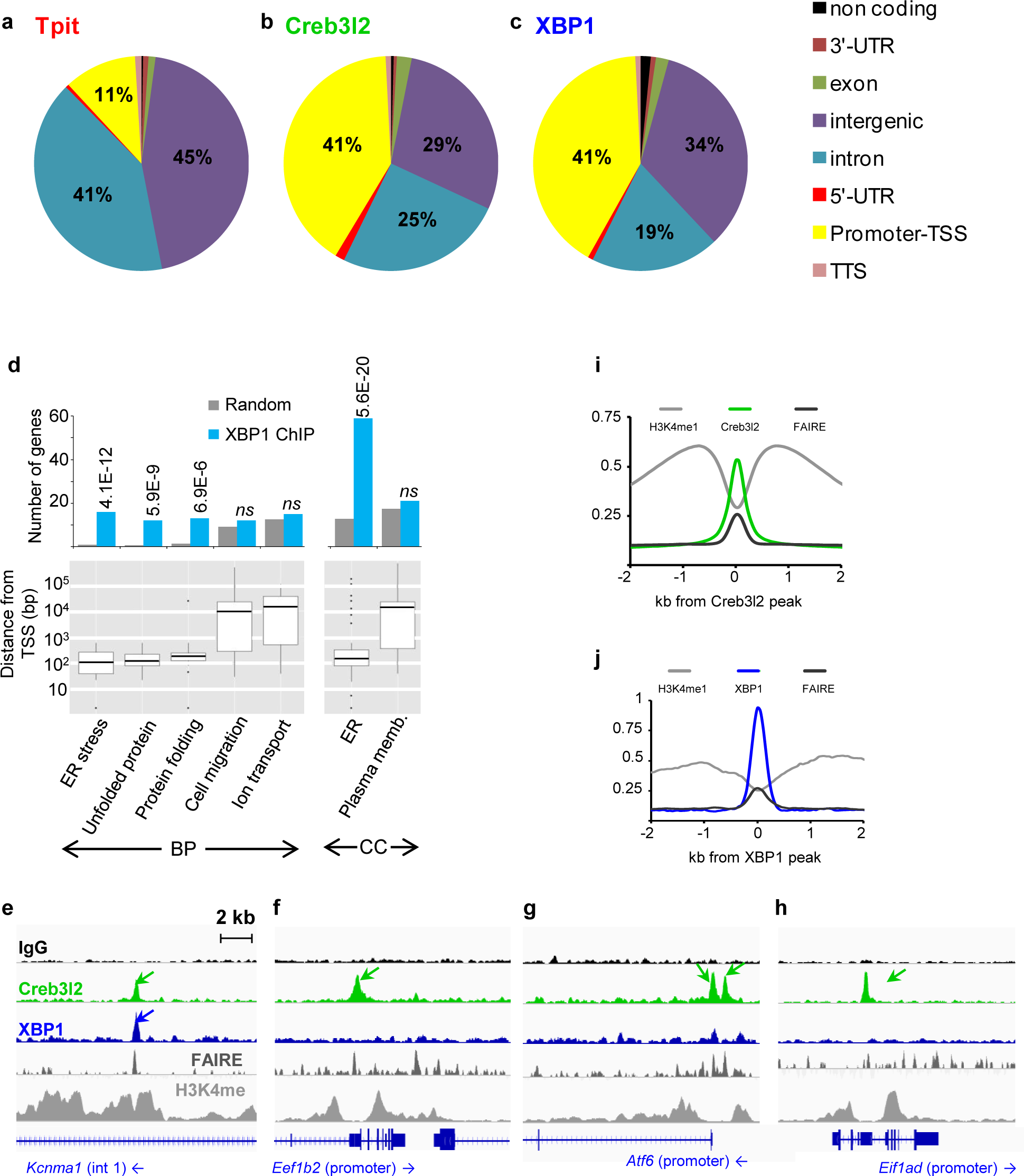
Creb3l2 and XBP1 target the promoters of translation and ER biogenesis genes, respectively. (**a-c**) Genomic distribution of Tpit, Creb3l2 and XBP1 ChIPseq peaks. (**d**) GO terms of genes associated with TSS proximal (≤1 kb) XBP1 peaks. Peaks were assigned to the closest gene with the AnnotatePeaks Homer command. Bars represent the number of genes in each category associated with XBP1 peaks (blue) or random occurrence of genes in each category (grey). P-values relative to random occurrence are shown above the bars; ns, not significant. The bottom panels provide box-plot representation of the distance to TSS for XBP1 peaks of each GO category revealing promoter-proximal associations for groups with significant associations. BP: biological process, CC: cellular component. (**e-h**) ChIPseq profiles at regulatory sequences of genes targeted by Creb3l2 and/or XBP1. ChIPseq patterns are shown for control IgG, Creb3l2, XBP1, H3K4me1 and FAIREseq. (**i-j**) Average profiles of H3K4me1 ChIPseq and FAIREseq at Creb3l2 (**i**) or XBP1 (**j**) peaks.

**Figure S4.**
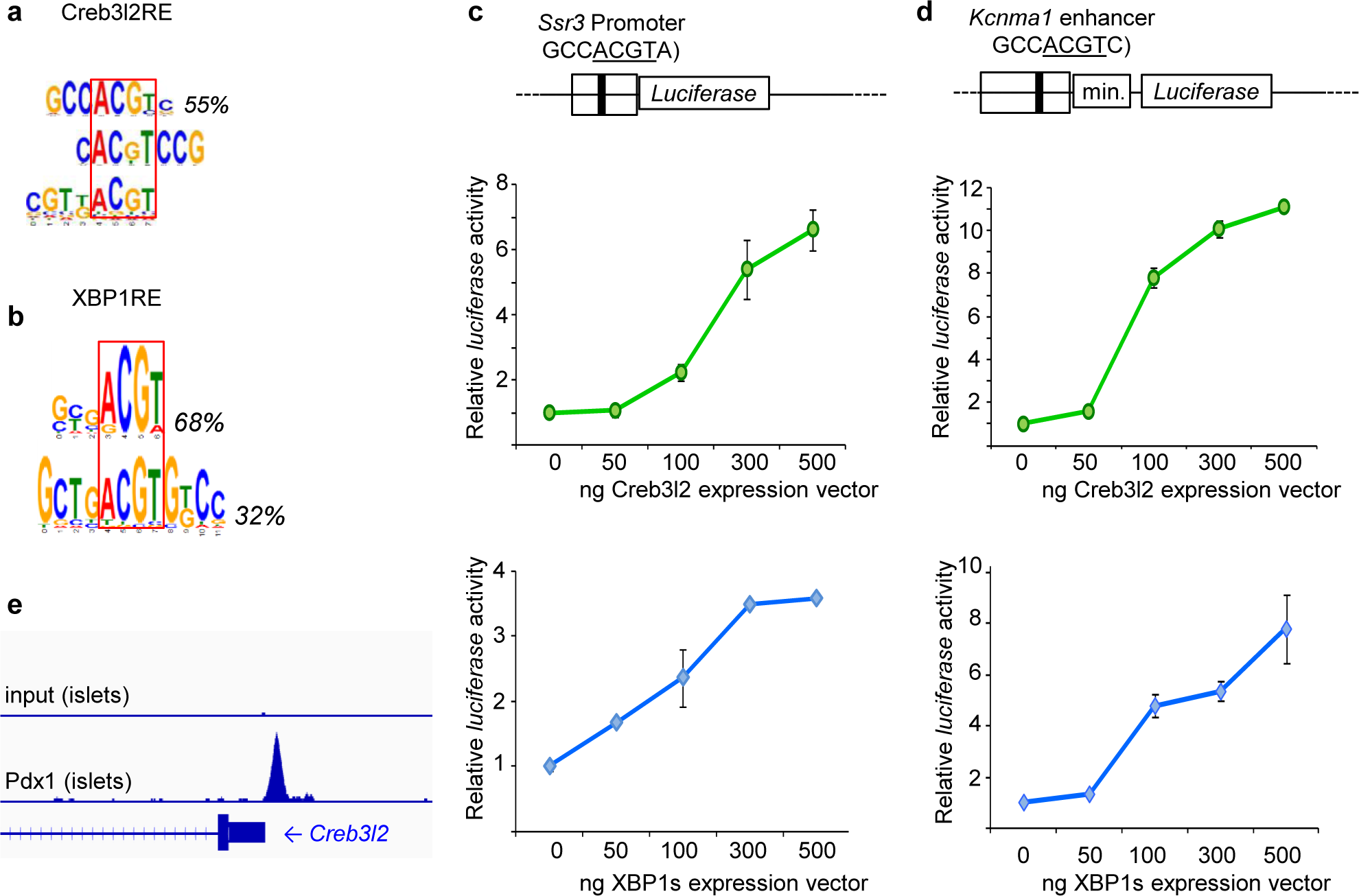
Creb3l2 and XBP1 activate transcription through binding of a similar sequence motif. (**a-b**) Creb3l2-RE and XBP1-RE consensus identified by *de novo* motif searches in sequences bound by Creb3l2 and XBP1. (**c-d**) The transcriptional activity of Creb3l2 and XBP1 assessed by transfection in GH3 cells using two pGL4.10-*luciferase* reporters. Schematic representation of *Luciferase* reporters containing the *Ssr3* promoter (**c**) or *Kcnma1* enhancer (**d**) and dose-response curves. (**e**) ChIPseq profiles of Pdx1 at regulatory sequences of Creb3l2 in pancreatic beta cells. Data from http://chip-atlas.org (the Pdx cistrome of pancreatic islets, ERX103428, ERX103429).

